# Contrasting roles of landscape compositions in shaping functional traits of arthropod community in subtropical vegetable fields

**DOI:** 10.1101/2022.07.13.499842

**Authors:** Jie Zhang, Hafiz Sohaib Ahmed Saqib, Dongsheng Niu, Karla Giovana Guaman Gavilanez, Ao Wang, Deyi Yu, Minsheng You, Gabor Pozsgai, Shijun You

**Affiliations:** State Key Laboratory of Ecological Pest Control for Fujian and Taiwan Crops, Institute of Applied Ecology, Fujian Agriculture and Forestry University, Fuzhou 350002, China; Fujian Key Laboratory for Monitoring and Integrated Management of Crop Pests, Institute of Plant Protection, Fujian Academy of Agricultural Sciences, Fuzhou 350013, China; Joint International Research Laboratory of Ecological Pest Control, Ministry of Education, Fuzhou 350002, China; Guangdong Provincial Key Laboratory of Marine Biology, College of Science, Shantou University, Shantou 515063, China; Ministerial and Provincial Joint Innovation Centre for Safety Production of Cross-Strait Crops, Fujian Agriculture and Forestry University, Fuzhou 350002, China; Ce3C - Centre for Ecology, Evolution and Environmental Changes, Azorean Biodiversity Group, CHANGE – Global Change and Sustainability Institute, University of the Azores, Faculty of Agricultural Sciences and Environment, Rua Capitão João D’ Ávila, São Pedro, 9700-042, Angra do Heroísmo, Terceira, Açores, Portugal; BGI-Sanya, Sanya 572025, China

**Keywords:** habitat transformation, biological pest control, functional diversity, trait composition, scale-dependent, RLQ analyses

## Abstract

Agricultural intensification and land use transformation are among the main driving forces of the unprecedented decline of biodiversity and ecosystem services in croplands. Trait-based approaches provide a unique framework to detect the potential mechanisms of how this intensification affects biodiversity and alter ecosystem services. However, the potential relationship between arthropod traits and various types of habitats is still poorly understood, especially in subtropical vegetable agroecosystems.

Here, we conducted a trait-based approach to evaluate the variable roles of different habitats on functional trait diversity and the structure of the arthropod community in brassica vegetable crops. Twenty-three conventional cruciferous vegetables fields were sampled over two years in three regions in Fujian, China. We found that the increasing proportion of non-brassica vegetable plantations and water bodies negatively affected the functional diversity of arthropods, whereas forest and grassland habitats showed a positive correlation, indicating habitat filtering for certain traits or trait combinations.

This study demonstrates the importance of landscape composition as an ecological filter for vegetable arthropod community, and identifies how the proportion of different habitats selected for or against specific functional traits. Our findings support that increasing forest and grassland areas adjacent to vegetable fields can play a vital role in promoting the functional diversity of arthropod communities. Since the natural enemy assemblages supported by these habitats bear combinations of diverse traits adapted to disturbance, they have the potential to enhance pest suppression in the highly variable environment of vegetable crops.

## 1. Introduction

Insect diversity is in a worldwide decline with insecticide spraying, natural habitats loss, and land use transformation contributing the most to the process (Didham et al., 2020). Biodiversity is also dramatically decreasing in agricultural landscapes and this decrease is mostly driven by enlarged field size and the intensification of management (Batáry et al., 2020). Consequently, the delivery of insect-related ecosystem services in agroecosystems, such as biological control, has also been hampered in the past decades (Martin et al., 2019b). Therefore, conservation biological control was proposed to increase the diversity of natural enemies, mostly by increasing landscape heterogeneity (Jonsson et al. 2008). Landscape heterogeneity, including landscape composition (diversity of habitat types) and landscape configuration (size, shape, and connectivity of habitats), is regarded as a crucial driver of biodiversity in agricultural landscapes (Fahrig et al. 2011). Studies have shown that diverse landscape compositions, such as the mixture of forest or grassland areas, are crucial for promoting natural enemies of pests by providing essential resources, such as pollen, nectar, and shelter (Gurr et al. 2017). Thus, increasing the proportion of seminatural habitats in the landscape is associated with the increase in the diversity of natural enemies, such as predators and parasitoids, which, in turn, provide a high level of pest control (Grab et al., 2018; Holland et al., 2016). However, some contrasting effects of landscape composition on pest control have been found across different agroecosystems and for different functional groups, sometimes positive and sometimes negative in landscapes with more noncrop habitat (Tscharntke et al. 2016; Karp et al. 2018). Therefore, the consensus is still lacking on the importance of landscape composition for beneficial arthropods and the ecosystem services they provide in crop fields.

Furthermore, recent developments in ecology suggest that taxonomic diversity tells us relatively little about the patterns of species assemblages driving specific ecosystem services (such as pest control) in agroecosystem (de Bello et al., 2010; Mayfield et al., 2010). Trait-based approaches, on the other hand, are thought to be superior to taxonomic indices in linking diversity to ecosystem functions (Gagic et al., 2015). This approach provides a unique framework to understand how environmental variables affect the function of arthropod communities (Mammola et al., 2021; Moretti et al., 2017; Wood et al., 2015). Investigating trait distributions in communities also can pinpoint how different habitats mosaics and agricultural intensification affect biodiversity and alter ecosystem services in agroecosystems (Fournier et al. 2015; Perović et al. 2018). For instance, studies have shown that small species with a shorter life cycle and higher dispersal ability have the advantage of avoiding frequent disturbance, and, as a consequence, these traits, highly adapted to the disturbed environment, were selected for in intensively managed grasslands (Simons et al., 2016). In contrast, more diverse trait communities were found in restored grasslands, as an outcome proving the success of restoration measures in herbivorous insect communities in seminatural grasslands (Neff et al., 2020). The effects of landscape heterogeneity and in-field management on the functional community composition of arthropods are relatively well-studied in temperate managed grassland ecosystems (Gámez-Virués et al. 2015; Simons et al. 2016; Neff et al. 2020) and tropical rice agroecosystems (Dominik et al., 2022). However, trait-based approaches on investigating cropland arthropod communities along landscape gradients have only recently received substantial research attention (Perović et al. 2018; Martin et al. 2019; Neff et al. 2020) and the insight into the roles various habitats play in shaping the trait composition in subtropical vegetable agroecosystems community is still rare (but see Saqib et al. (2022) for trait compositions of spider assemblages in vegetable fields).

Moreover, the impact of functional diversities on ecosystem services, as well as the landscape processes driving functional diversities, are most likely scale-dependent (Suárez-Castro et al. 2022; Wood et al., 2015) and neglecting this scale dependency may hamper the effective conservation biological control in agriculture landscapes. Yet, our understanding of the effects of seminatural habitats on arthropod functional diversity at multiple spatial scales is largely limited. Indeed, other than a few examples, the majority of the studies on the effects of functional diversity on ecosystem functioning were conducted on a fine (and/or a single) spatial scale (Gonzalez et al., 2020). Although Zhang et al. (2021) showed varying effects of habitats on the taxonomic diversity of arthropod communities across five different spatial scales in managed brassica vegetables, it is largely unknown if these changes in taxonomic diversity are accompanied by similar trends in functional diversity.

In order to address both the landscape- and scale-dependency of arthropod trait diversities in agroecosystems, in this study, we evaluated the effects of habitats on the functional diversity and composition of arthropod communities in cruciferous vegetables at five different spatial scales. We sampled 23 brassica fields over a gradient of seminatural habitats (e.g. forests and grasslands) in the adjacent landscape mosaic, and assessed the functional assemblages of both pests and natural enemies over two years in three typical cruciferous vegetable planting regions in Fujian, southeast China. We hypothesized that:

(1) Increasing proportion of seminatural habitats (e.g. forest and grassland) is in positive correlation with trait diversity of the arthropod community, but managed habitats (e.g. vegetables) negatively affect them.
(2) Various habitats filter specific traits of the arthropod community.
(3) Similarly to taxonomic diversity patterns, the response of functional diversities to different habitats varies across scales.

## 2. Material and Methods

### 2.1 Study area and land use analysis

The study was carried out in Fujian, southeast China (Figure 1a). In total, 23 fields were selected along a gradient of proportion of seminatural habitats, all fields were at least 1 km apart from each other, at three sampling regions, situated in conventional vegetable planting areas: (1) a vegetable-rice rotation cropping system in the hilly lowlands of Nanping Municipal Region in northwest Fujian defined by agroforestry with narrow plains between river and mountain; (2) an intensively cultivated vegetable-rice rotation cropping system in the coastal plain of Zhangzhou Municipal Region in southern Fujian characterized by large vegetable monocultures and few seminatural habitats; and (3) a traditional multi-type vegetable continuous cropping system in the mountainous Ningde Municipal Region located in the northeast of Fujian where small patches of vegetable habitats and large patches of forest dominate the landscape.

**Figure 1.**
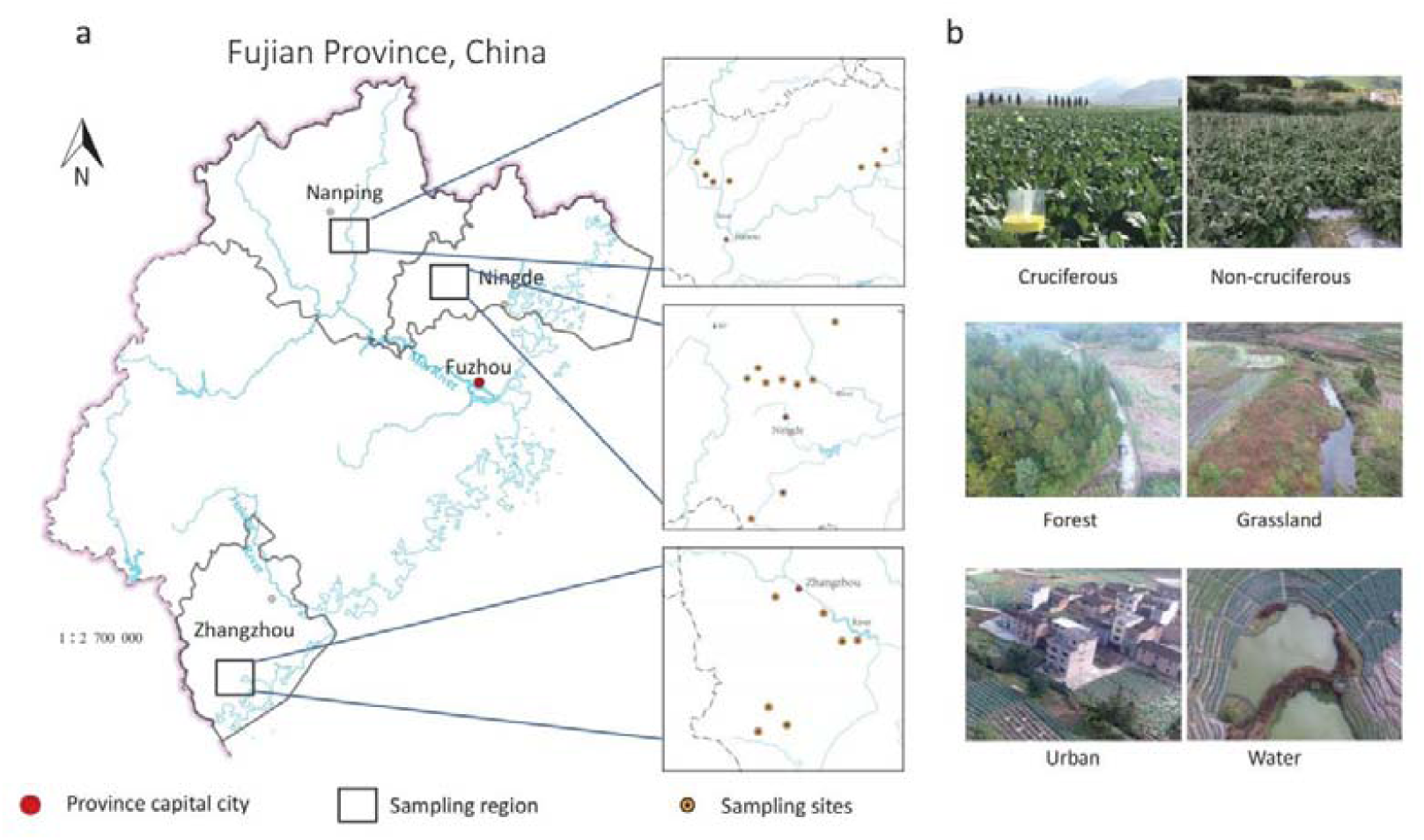
Map showing (a) sampling regions in Fujian, southeastern China, and (b) examples of different habitats.

To investigate the land use distribution in each field, aerial photos were taken by a drone (PHANTOM 4, Shenzhen Dajiang Baiwang Technology Co., Ltd., China), then data were validated by field investigations. Land uses were categorized into one of the six types: cruciferous vegetable (cauliflower) plantation, non-cruciferous vegetable (e.g. eggplant, pepper, and beans) plantation, grassland, forest, urban areas (e.g. residential lands or roads), water surfaces (e.g. small streams or ponds) (Figure 1b). The proportion of different land use types was calculated for five different spatial scales, 100, 200, 300, 400, and 500 meter radius around the focal field, by using QGIS 3.4 (for details, see Zhang et al. (2021)).

### 2.2 Arthropod sampling and identification

Sampling was conducted in the main cauliflower growing season (from April to May) for two years (2017 and 2018). Arthropods were sampled three times during the pre-harvest period because it is generally associated with a maximum abundance of arthropods. Airborne arthropods (e.g. aphids, Vespidae wasps, flies, ballooning spiders, and parasitic wasps) were sampled by yellow pan traps. Five pan traps at vegetable height were set in each focal patch (one in the center and four in the corner of the square with 20 m apart from each other). Ground-dwelling arthropods (e.g. spiders, carabids, and rove beetles) were sampled using pitfall traps (9.5 cm diameter and 13.8 cm deep) with a cover above each pitfall trap to prevent flooding by rainwater. The pitfall traps were half-filled with propylene glycol solution with a drop of added detergent to reduce surface tension. They were set at each site in the same pattern as the pan traps. Besides pitfall traps, boat-shape traps with a sex pheromone lure were set to sample *Plutella xylostella* (Linnaeus, 1758) males (Zhang et al., 2021). All specimens were transferred into 80% ethanol in tubes and identified to the lowest possible taxonomic level using the molecular barcoding procedures given by Saqib et al. (2022). Although species-level identification was not always possible, even family-level identifications have been proved to provide ecologically adequate surrogates for species in the study of functional diversity (Lamarre et al., 2016). Thus, species whose species identity was not possible to resolve were included in the analysis at the family-level as unknown species.

### 2.3 Trait definition and measurement

Information of seven functional traits (feeding mode, diet breadth, agricultural specialism, overwintering habitat, dispersal mode, stratum, and activity period) was extracted from the published scientific literature for all species included in the study (Table 1). These traits were selected because of their potential influence on arthropod responses to landscape variables through varying ways of movement or dispersal between different habitats, getting food, finding hosts, or avoiding disturbance (Martin et al., 2019a).

**Table 1.**
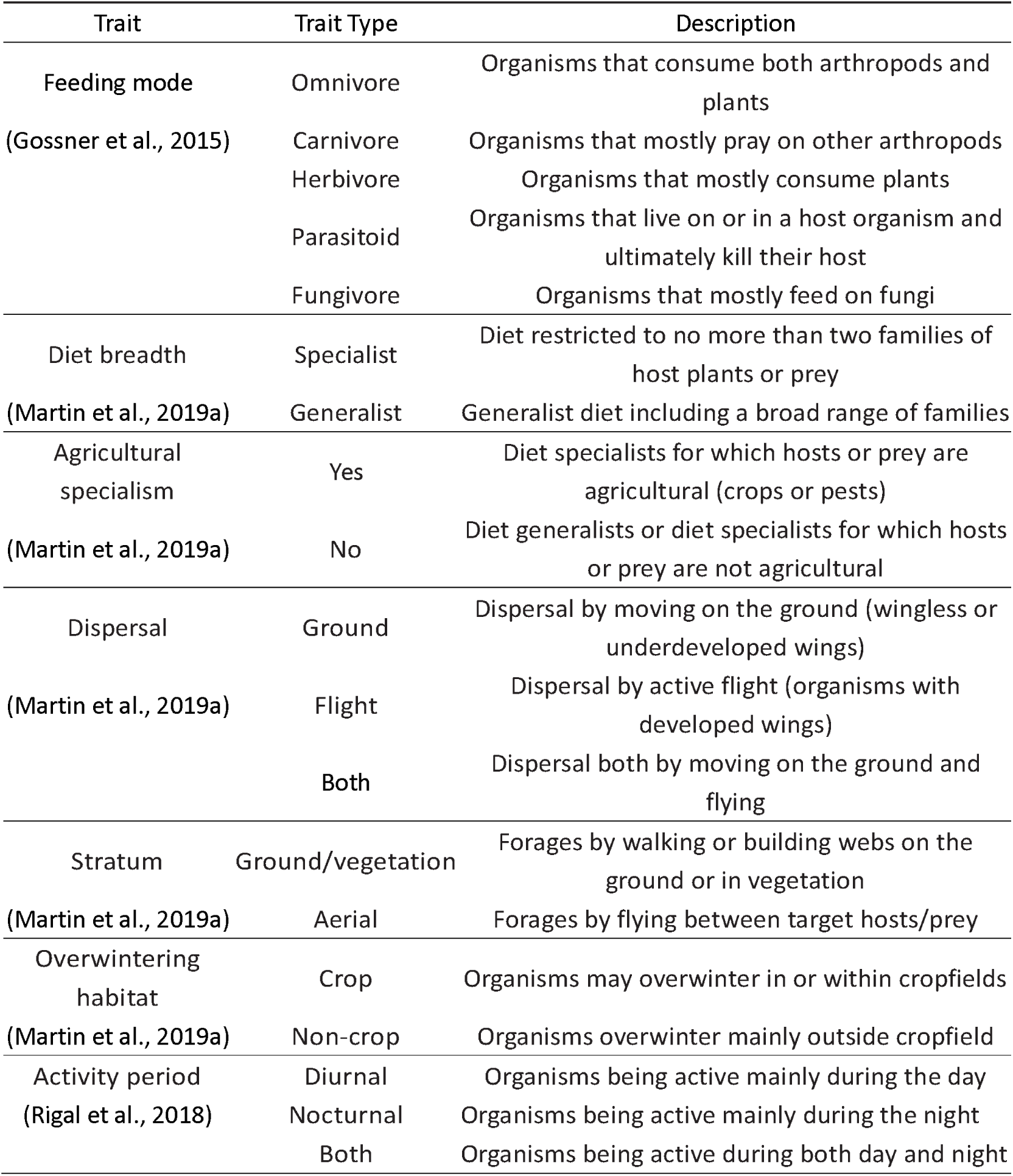
Functional traits of arthropods that related to habitats.

### 2.4 Statistical analysis

#### Functional trait diversity

To investigate how the functional trait diversity of arthropod communities is associated with different types of habitats in a vegetable growing landscape mosaic at varying spatial scales, we first computed the three most widely used functional diversity indices in each field: functional richness (FRic) (Villéger et al., 2008), functional dispersion (FDis) (Laliberte and Legendre, 2010) and functional evenness (FEve) (Botta-Dukát, 2005), using the function *dbFD*() from the FD package in R (Laliberté et al., 2014).

We then used a series of generalized linear mixed-effects models (GLMM) to analyze the relationship between the functional diversity and landscape composition. For the individual models, (1) FRic, (2) FEve, and (3) FDis were used as response variables, and the proportion of all five habitat types at each field as explanatory landscape composition variables. Sampling regions were included as interacting variables with landscape composition variables. Study field and sampling year were added as random variables. To check for multicollinearity between explanatory variables, variance inflation factor (VIF) was calculated by the “car” R package (Fox and Weisberg, 2019). Cruciferous vegetables were excluded in our final models for its high collinearity with other habitat proportions (VIF > 5) (Zuur et al., 2009). Analyzes were conducted at all five spatial scales. All predictor variables were standardized (z-scored transformation) prior to analysis (Grueber et al., 2011) using the *standardize*() function of the R package “arm” (v. 1.11-2; Gelman et al., 2020). All possible combinations of models were fitted and their associated corrected Akaike’s Information Criteria (AICc) were calculated (Burnham et al., 2011) using the *dredge*() function of R package “MuMin” (v. 1.43.1; Barton, 2020). The best fitted models were obtained (ΔAICc < 2) with the *get*.*models*() function and averaged using the *model*.*avg*() function of package “MuMin”.

#### Functional trait composition

To test the individual trait-environment relationship, we used a combination of the RLQ method with the fourth-corner approach as described by Dray et al. (2014). This approach uses two complementary multivariate analyses linking data from three tables: environmental variables per site (R), species relative abundances per site (L), and functional traits per species (Q), which allows to better explore multivariate patterns and test for the significance of multivariate associations. Prior to the RLQ analyses, a Hellinger transformation (Legendre and Gallagher, 2001), as implemented in the “vegan” package in R (Oksanen et al., 2019), was applied to standardize the species abundance matrix. Then the L species abundance table was analyzed using correspondence analysis (CA), and the Q species trait and R environmental tables were analyzed by the Hill-Smith principal components analysis (PCA) (Hill and Smith, 1976). Monte-Carlo testing was applied to detect the statistical significance of relationship among three RLQ tables using the *fourthcorner2*() function in “ade4” package (Dray and Anne, 2007). Then the statistical significance of individual trait-environmental relationships was evaluated by the fourth-corner test with 49,999 permutations using the *fourthcorner*.*rlq*() function. Considering the multiple tests on environmental variables, P-values were adjusted by the false discovery rate method (FDR; Benjamini and Hochberg 1995; Dray et al. 2014). Then clusters were identified to separate the responsive and non-responsive traits based on Euclidean distances along the first two axes, using Ward’s hierarchical clustering method. Then Calinski–Harabasz stopping criterion (Calinski and Harabasz, 1974) was applied to determine the optimal number of functional groups. The clustering of RLQ analysis results was performed following Kleyer et al. (2012).

We had tested the spatial autocorrelation in the co-inertia remained unexplained for each of the first two RLQ axes using *gearymoran*() function in “ade4” package, and spatial autocorrelation was not observed. All analyses were performed using R 3.6.2 (R Core Team 2019).

## 3. Results

### 3.1 Effects of habitats on functional diversity

We collected 137,726 arthropod individuals from 127 families, 15 orders (Table S1) in total. Results show that the proportion of forest in the landscape was positively associated with FRic at all investigated spatial scales, and that the proportion of grassland was positively associated with FDis and FEve at larger scales (300 - 500m) but not with FRic. Conversely, the proportion of the area covered with non-cruciferous vegetables and water negatively correlated with FEve at 100m and 500m, respectively (Figure 2, Table S2). Besides, sampling regions also significantly affected trait diversity with FRic and FEve being markedly lower in Zhangzhou region than in the other two regions (Figure 3, Table S2).

**Figure 2.**
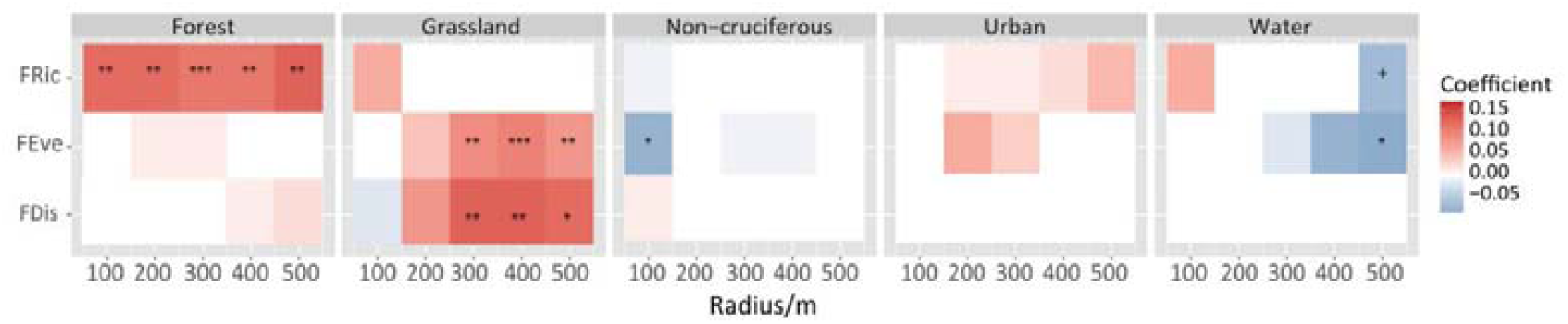
Relationship between the functional diversity index (FRic, FEve and FDis) and landscape composition variables at multiple spatial scales. The gradient of the color represents the value of the GLMM estimated coefficients. “*” indicates a significant correlation at P < 0.05 level, “**”at P < 0.01 level, and “***”at P < 0.001 level. “+” indicates the interaction between landscape variables with sampling regions.

**Figure 3.**
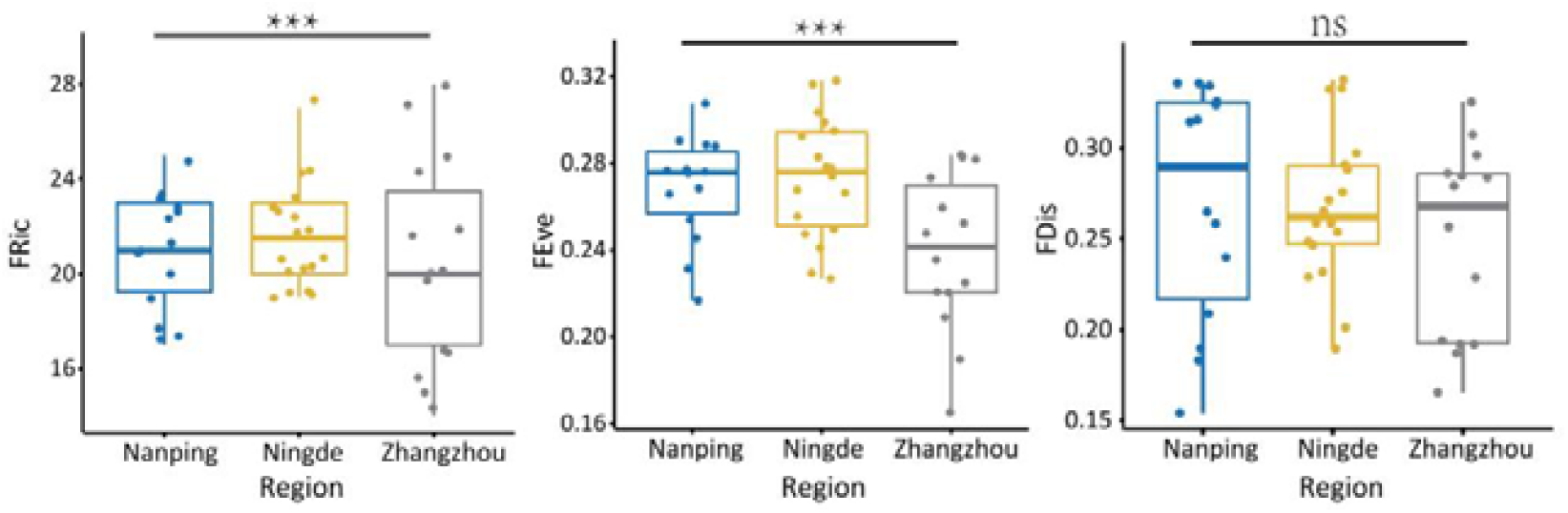
Functional diversity (FRic, FEve and FDis) in different sampling regions. “***” represents significant effects (P < 0.001) of sampling regions measured by the GLMM test. “ns”indicates there is no significant correlations between sampling regions.

### 3.2 Effects of habitats on functional composition

The Monte-Carlo tests showed a significant relationship (p < 0.001) between the three RLQ tables (including species abundance, functional traits and habitat variables). The first two axes of the RLQ analyses explained 85% of total co-inertia between arthropod functional traits and environmental variables (including sampling region, radius and proportion of different habitats).

The high proportion of grasslands was associated with non-agricultural specialism, including carnivore and fungivore species (i.e. members of the Theridiidae spider, and Carabidae and Cryptophagidae beetle families, Figure 4), with nocturnal activity, overwintering mainly outside crop fields, and dispersal mode both by flying and on the ground in Cluster B (Figure 5, Figure 6);

**Figure 4.**
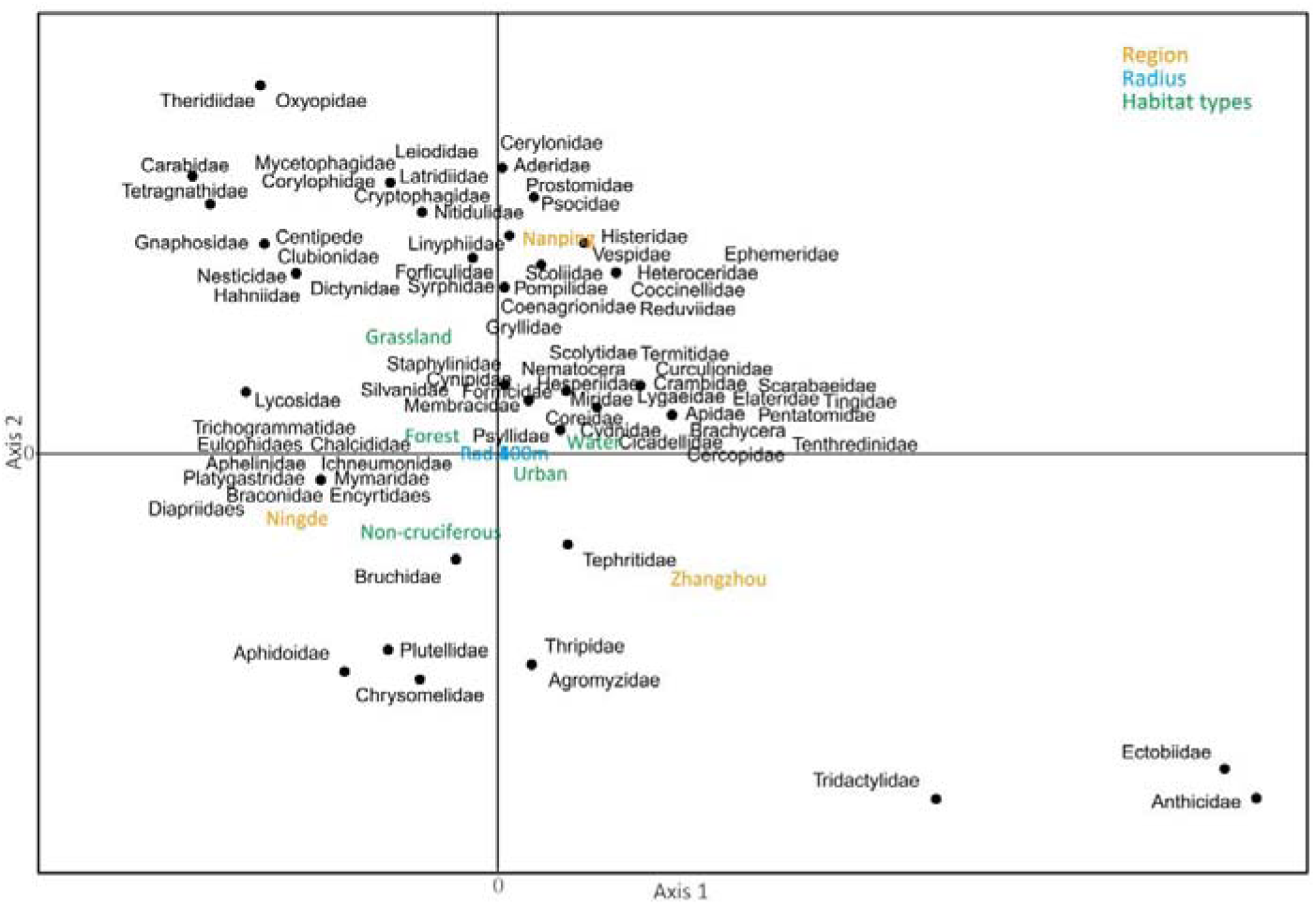
The association of sampling regions, landscape spatial scales, proportion of different habitats in surrounding landscape with families of arthropods along the first two axes of the RLQ analysis. Each dot represents a different family, individual number >10.

**Figure 5.**
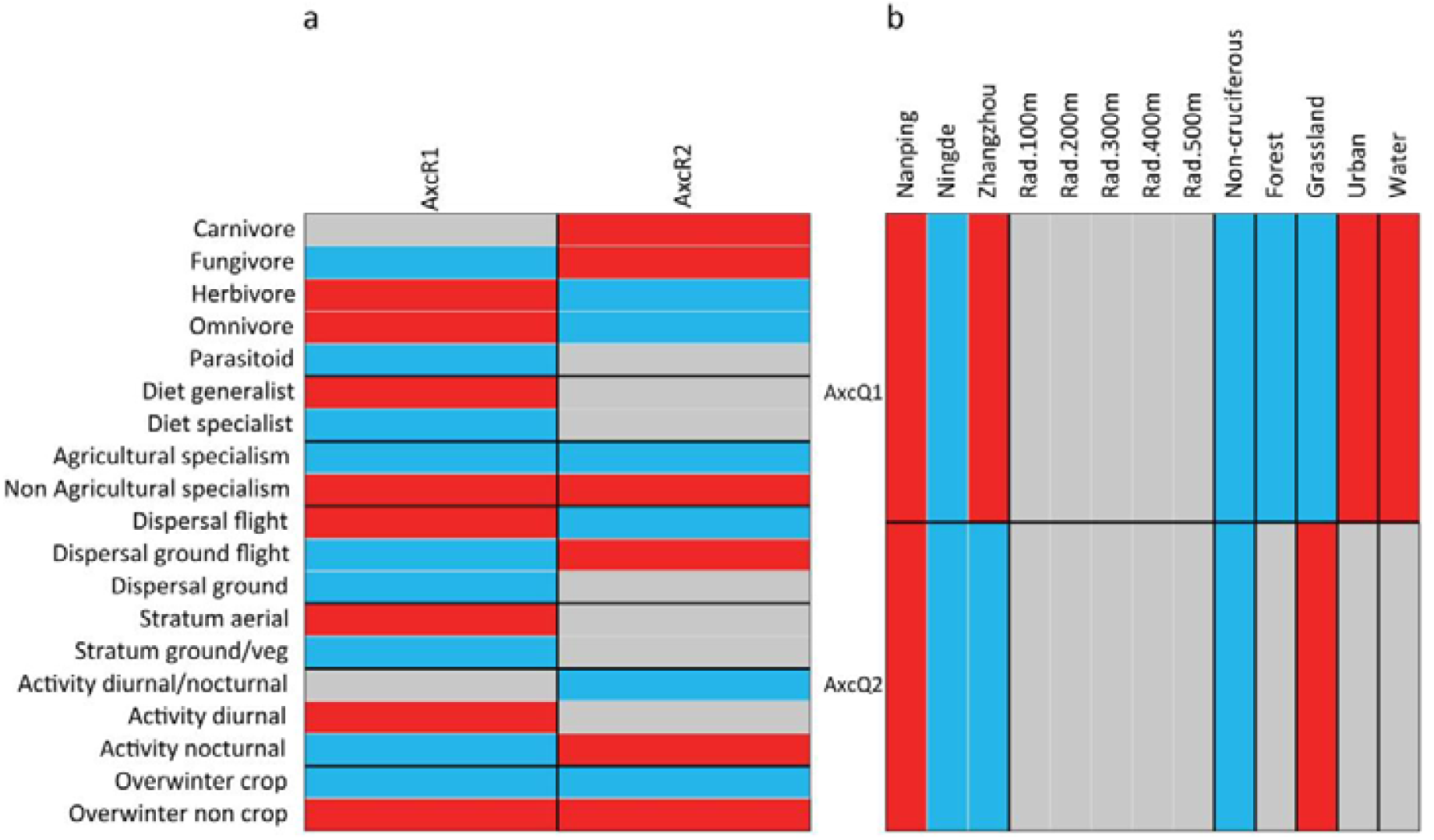
Combination of the fourth-corner and RLQ results. (a) Fourth-corner tests between the first two RLQ axes for landscape gradients (AxcR1/AxcR2) and functional traits, and (b) fourth-corner tests between the first two axes for functional traits (AxcQ1/AxcQ2) and landscape variables. Red cells represent positive significant (*P* < 0.05) associations, and blue cells significant negative associations. Non-significant (*P* > 0.05) associations are shown in gray cells.

**Figure 6.**
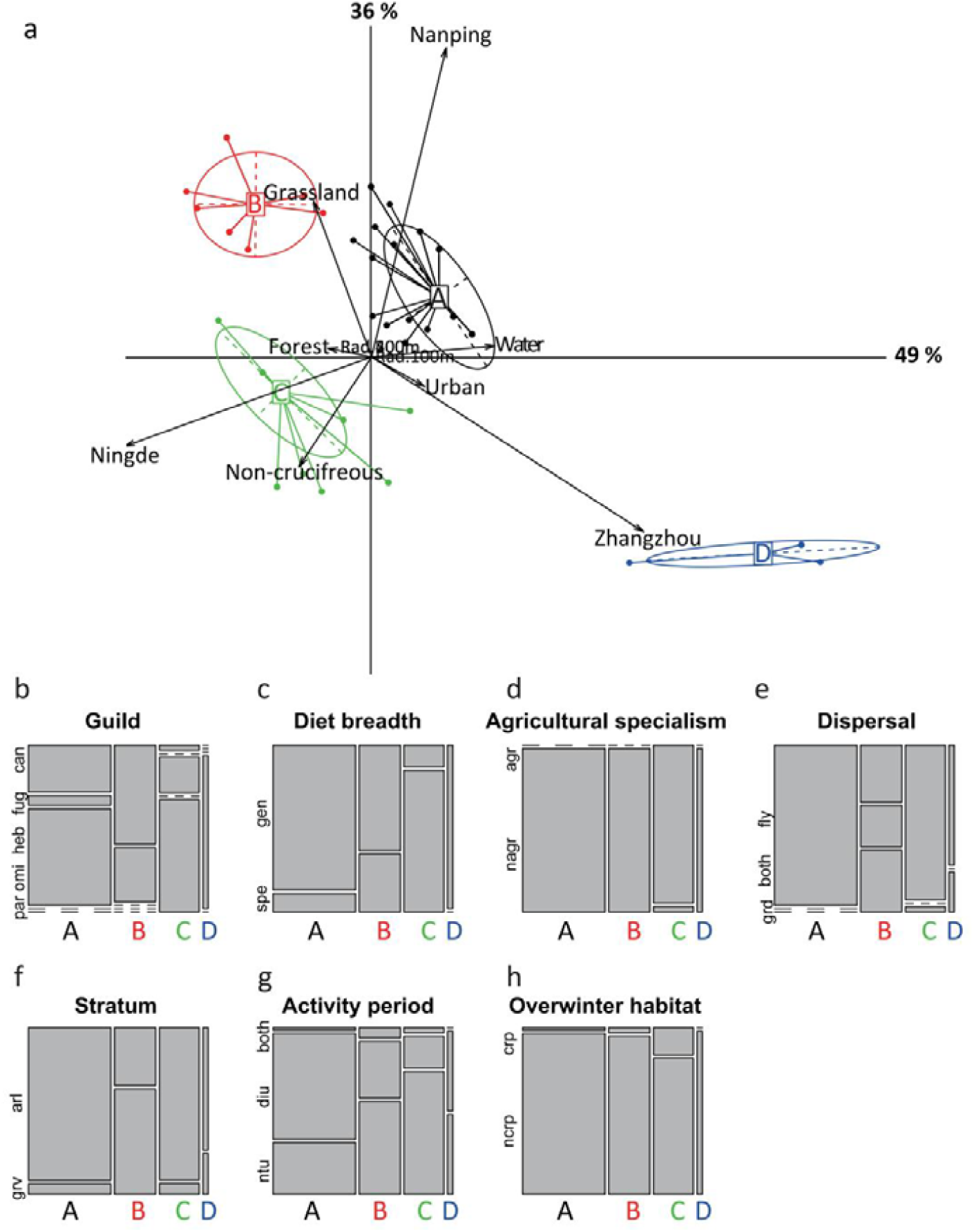
The association of sampling regions, landscape spatial scales, proportion of different habitats in surrounding landscape with traits classification of arthropods. (a) RLQ biplot, showing the clustering of co-associations between habitats variables and functional trait attributes, weighted by abundance. Arrows represent the size and direction of landscape factors effects. Clustered points show the position of derived functional groups and are represented with the same color (“A”, “B”, “C” and “D”). (b-h) Rectangle bars represent the distribution of functional trait attributes in cluster “A”, “B”, “C” and “D”. The width of the rectangle bars represents relative abundance, and the height represents the proportion of different functional attributes within cluster “A”, “B”, “C” and “D”: the higher the bar, the greater the proportion functional attribute. (can-carnivore, fug-fungivore, heb-herbivore, omi-omnivore, par-parasitoid; spe-specialist, gen-generlist; agr-agriculture, nagr-non agriculture; grd-ground, fly-fly; arl-aerial, grv-ground/vegetation; ntu-nocturnal, diu-diurnal; crp-crop, ncrp-non crop).

The high proportion of forest in the landscape supported agricultural specialism and overwintering mainly within crop fields, as well as specialist Hymenoptera parasitoids (i.e. Braconidae, Ichneumonidae, and Encyrtidae, Figure 4) which were characterized by nocturnal activity, dispersing by both flying and on the ground, foraging both on the ground and vegetation in Cluster C (Figure 5, Figure 6). The increasing proportion of non-cruciferous crop growing areas (mainly eggplant, pepper and soybean) supported the same Cluster C as the forest area proportion (Figure 6a), and, similarly, the community mainly consisted of agricultural specialists. However, in the latter case, they were more herbivores (i.e. Lepidoptera: Plutellidae, Hemiptera: Aphidoidae and Diptera: Agromyzidae, Figure 4), with both diurnal and nocturnal activity and with a flying dispersal type (Figure 5).

Both water surfaces and urban areas supported non-agricultural generalists, with diurnal activity, flying dispersal, aerial foraging, and overwintering outside of crop fields (Figure 5). Unlike urban habitats, which mainly supported more omnivores in Cluster D (ie, Tridactylidae, Ectobiidae and Anthicidae, Figure 4), water surfaces supported herbivores (ie, Lygaeidae, Miridae and Apidae, Figure 4) in Cluster A (Figure 6a and Figure 6b).

## 4. Discussion

In this study, we analyzed the roles of different habitats in forming functional trait diversity patterns and filtering trait composition of taxonomically diverse arthropod communities along five spatial scales in brassica vegetable plantations. We found that habitats surrounding vegetable plantations influenced arthropod functional diversity in the field, but different habitats played contrasting roles in filtering for and against trait combinations.

The functional richness of the arthropod community in vegetable growing areas was markedly supported by the surrounding forest habitats but not by non-brassica crop areas. Since forests or woodlands in the subtropics can provide a stable environment with a continuous supply of resources for various arthropods, and thus help them to avoid spatial and temporal disturbances (Diekötter and Crist, 2013; Schoenly et al., 2010), these habitat patches may act as a source of diverse arthropod communities on brassica fields. Indeed, increasing forest proportions supported broad trait spaces (e.g. dispersal or forages by flying and ground, forages on the ground and vegetation), and protected those traits that are not adapted to human disturbance (e.g. agricultural specialism, specialist organisms). In our study, the agricultural specialists supported by forests were mainly parasitoids, such as Braconidae and Ichneumonidae wasps, which can travel great distances for searching for hosts. This agrees with previous studies which showed that arthropods specialized in their feeding were favored by the greater diversity of land cover types (Gámez-Virués et al., 2015) which would be lost with increasing land-use intensity (Simons et al. 2016). Although studies found no evidence that landscape composition positively affects the abundance of parasitoids in rice agroecosystem (Sann et al., 2018; Zou et al., 2020), our results suggested that increasing forest area in the vegetable landscape mosaic supported the parasitoid community, especially Braconidae, Ichneumonidae, Encyrtide and Eulophidae wasps. Apparently, forest area positively affected functional richness and supported broad trait space. Thus, to a certain extent, our results support the landscape-moderated insurance hypothesis, that forest area provides insurance (resilience and stability) for functional traits when those face the adverse effects of intense management (Tscharntke et al. 2012; Gámez-Virués et al. 2015).

Our results showed that the increasing proportion of grassland area also increased functional evenness and functional dispersion; two measures indicating the resistance and stability potential of communities as well as their capability to offset the negative effects of management (Mason et al. 2005). The positive relationship with functional dispersion in our study indicates that increasing proportion of grasslands could maintain a arthropod communities with a high degree of niche differentiation and low level of resource competition (Mason et al., 2005), whilst high functional evenness indicates low habitat dynamics and efficiently utilized niche space, which tend to increase productivity and reliability (Mason et al., 2005; Schleuter et al., 2010). That non-agricultural specialist species, especially carnivores (e.g. ballooning spiders and carabid beetles) which are able to disperse both by flying and on the ground were largely promoted by grasslands further underlines the assumption that grasslands can serve as little disturbed, highly productive habitats, with low competition, which help functional groups to spill over into cauliflower. Indeed, in agricultural landscapes, the majority of grasslands are situated along the field edges and less likely disturbed than planting areas. They play a prominent role in shaping in-field trait compositions of the arthropod community, including those normally considered natural enemies, such as spiders and carabids (Schirmel et al. 2016; Gallé et al. 2020; Saqib et al. 2022), which overwinter mainly outside the crop and whose nocturnal activity helps them to avoid frequent human disturbance. Thus, increasing the proportion of grasslands has the merit of improving or maintaining the community resistance to artificial disturbance by improving niche complementarity and resource utilization (Prada-Salcedo et al., 2021).

Conversely, the proportion of non-cruciferous crop plantations showed markedly negative effects on functional evenness of the arthropod community. Since these habitats are generally associated with higher chemical inputs (i.e., pesticides, fertilizers, etc.), only species with high competitive ability and stress tolerance are likely to survive and become dominant (Mayfield and Levine, 2010). Thus, by filtering to similar traits, this process leads to lower functional evenness. Especially, herbivore species that are agricultural specialists with high dispersal ability (e.g. dispersal by flying) were selected under intensive management. These results corroborate the findings of Birkhofer et al. (2015) who showed that the more frequently disturbed land-use types were characterized by species with higher dispersal abilities and communities with lower functional distinctness or trait richness. Moreover, communities with a broad temporal niche (activity of both diurnal and nocturnal) were common in landscapes with a high proportion of non-cruciferous crops. The longer activity periods could help these arthropods to fully utilize agricultural resources and maintain populations in intensively managed landscapes. This is supported by the finding that species with shorter activity periods are more likely to be absent in intensively managed agroecosystems (Gámez-Virués et al., 2015). In line with our results, both highly dispersive and broad temporal niche communities were enhanced in high-intensity land uses (Rigal et al., 2018).

Furthermore, consistent with our Hypothesis 3, and in line with the recent review of Suárez-Castro et al. (2022), the relationship between indices of functional diversity and landscape composition was highly variable across spatial scales. The response scales of FEve to grassland (300–500 m) and to water (500 m) were larger than to non-cruciferous crops (100 m). Although differences in the average dispersal ability between communities supported by different habitats may be one of the drivers, other life history traits such as diet breadth are also likely played a role. For instance, whereas abundant host plants at the 100m radius scale, where both cruciferous and non-cruciferous vegetables were the most common, supported agricultural herbivores (e.g. Plutellidae, Aphidoidae), landscapes more interwoven with grassland patches at larges scales supported non-agricultural carnivores (e.g. ballooning spiders and Carabidae), and water surfaces attracted both non-agricultural herbivores and carnivores (e.g. Syrphidae, Apidae and Reduviidae). Indeed, unevenly distributed landscape elements across spatial scales (Figure S1) can explain some of the patterns we found: within a 100 m radius from the central point of our sampling area, high intensity managements in non-cruciferous crops was the dominant landscape element and the best predictor of trait diversity indices. At a 300-500 m scale, the proportion of non-agricultural landscapes increased and the effects of more stable habitat conditions on trait diversities, mostly provided by grassland mosaics, became dominant. Although forests and woodlands were also represented in lower proportion at the 100 m scale, they equally positively affected FRic at all investigated spatial scales, indicating their remarkable capability of hosting diverse arthropod communities even small areas. Greater environmental heterogeneity supports higher levels of traits nestedness and turnover at landscape scales which lead to high FRic (Jarzyna and Jetz, 2018), and medium disturbance level is expected to increase FEve Suárez-Castro et al. (2022), the functional diversity indices are unlikely to respond to low-level habitat diversification in the very center of the agricultural field. Thus, to improve the efficiency of designing agricultural landscapes, at least a medium spatial scale (a radius of 300 m) of management is recommended.

Functional diversity indices were significantly influenced by sampling regions, yet no interacting effects were found with habitat proportions. The lowest functional richness and evenness were found in Zhangzhou region. In this area, large monoculture patches with intensive human disturbance are common which can be the main filtering factors that promote niche convergence, producing lower than expected functional richness and evenness in the communities. This is in line with a previous study that found ant assemblages with the lowest functional richness in simple landscapes (García-Navas et al., 2022) and lower functional evenness with increasing urbanization (Braschler et al., 2021). It also seems that regional differences are similarly crucial in shaping the functional diversity of arthropod communities as they are in shaping taxonomic diversity (Dominik et al., 2017; Massaloux et al., 2020).

## 5. Conclusions

In this study, we demonstrate the contrasting roles of different habitats as ecological filters of the functional trait diversity and trait composition of arthropod communities in vegetable fields at multiple spatial scales. Unlike agricultural land, which favors more herbivores with less specific traits adapted to managements, seminatural habitats promote carnivorous and parasitoid arthropods with higher functional diversity. Here, we provide evidence that the increasing proportion of seminatural habitats surrounding vegetable fields supports a functionally more diverse natural enemy communities (e.g. especially predators and parasitoids) with a broader foraging stratum and disparate ways of dispersal that are likely to increase the top-down control on all herbivore groups. To take full advantage of the biological control services, trait diversity maximization and trait composition optimization are recommended when designing agricultural landscapes. Specifically, seminatural habitats preserved at least 300 meters around vegetable fields are expected to maintain natural enemy communities with a high enough trait diversity for efficient biological control.

## Supporting information

able S2

## Authors’ contribution

JZ, GP, SY and MY conceived the ideas and design of the study and led the writing of the manuscript. JZ, DN, KGGG and AW conducted fieldwork. JZ, HSAS and GP analyzed the data. All authors revised the final version and gave their approval for submission.

## Acknowledgements

We thank Ibtissem Ben Fekih and David Perovic for their kind suggestions how to improve the manuscript. We thank all the land owners and farmers for their permissions and help which allowed us to sample their vegetable fields. We also thank Jun Li, Tao Li, and Lingfei Peng for their help in identifying the parasitoid samples.

## Funding

This work was financially supported by the National Natural Science Foundation of China (No. 31972271), the Fujian Agriculture and Forestry University Science and Technology Innovation Fund Project (CXZX2019001G, CXZX2017206), and the Outstanding Young Scientific Research Talents Program of Fujian Agriculture and Forestry University (xjq201905).

## Notes

### Competing Interest Statement

The authors have declared no competing interest.

