## Supplementary material for "Contrasting roles of landscape compositions in shaping functional traits of arthropod community in subtropical vegetable fields": Table S1

**Table S1 The list of the arthropod community and the functional traits(family level)**

(Forg-Foraging, DB-Diet breadth, AS-Agricultural specialism, DP-Dispersal, ST-Stratum, AP-Activity period, OH-Overwintering habitat)

|  | Family | | Order | | Forg | | DB | | AS | | DP | | ST | | AP | | OH | | 2017 | | 2018 | Sum | | |
| --- | --- | --- | --- | --- | --- | --- | --- | --- | --- | --- | --- | --- | --- | --- | --- | --- | --- | --- | --- | --- | --- | --- | --- | --- |
|  | Staphylinidae | | Coleoptera | | Carnivore | | Generalist | | No | | Flight | | Ground/Veg | | Diurnal | | Crop | | 1832 | | 2822 | 4654 | | |
|  | Linyphiidae | | Araneae | | Carnivore | | Generalist | | No | | Flight | | Ground/Veg | | Diurnal | | No | | 198 | | 366 | 564 | | |
|  | Carabidae | | Coleoptera | | Carnivore | | Generalist | | No | | Both | | Ground/Veg | | Nocturnal | | Crop | | 510 | | 492 | 1002 | | |
|  | Coccinellidae | | Coleoptera | | Carnivore | | Generalist | | No | | Flight | | Aerial | | Diurnal | | No | | 66 | | 51 | 117 | | |
|  | Forficulidae | | Dermaptera | | Carnivore | | Generalist | | No | | Flight | | Ground/Veg | | Nocturnal | | No | | 36 | | 78 | 114 | | |
|  | Formcidae | | Hymenoptera | | Carnivore | | Generalist | | No | | Flight | | Aerial | | Both | | No | | 65 | | 485 | 550 | | |
|  | Vespidae | | Hymenoptera | | Carnivore | | Generalist | | No | | Flight | | Aerial | | Diurnal | | No | | 17 | | 15 | 32 | | |
|  | Syrphidae | | Diptera | | Carnivore | | Specialist | | No | | Flight | | Aerial | | Diurnal | | No | | 302 | | 74 | 376 | | |
|  | Chrysididae | | Hymenoptera | | Carnivore | | Generalist | | No | | Flight | | Aerial | | Diurnal | | No | | 6 | | 1 | 7 | | |
|  | Chrysopidae | | Neuroptera | | Carnivore | | Generalist | | No | | Flight | | Aerial | | Nocturnal | | No | | 2 | | 5 | 7 | | |
|  | Reduviidae | | Hemiptera | | Carnivore | | Generalist | | No | | Flight | | Aerial | | Diurnal | | No | | 10 | | 10 | 20 | | |
|  | Aphelinidae | | Hymenoptera | | Parasitoid | | Specialist | | Yes | | Flight | | Aerial | | Nocturnal | | No | | 141 | | 186 | 327 | | |
|  | Bethylidae | | Hymenoptera | | Parasitoid | | Specialist | | Yes | | Flight | | Aerial | | Nocturnal | | No | | 9 | | 9 | 18 | | |
|  | Ceraphronidae | | Hymenoptera | | Parasitoid | | Specialist | | Yes | | Flight | | Aerial | | Nocturnal | | No | | 28 | | 209 | 237 | | |
|  | Chalcididae | | Hymenoptera | | Parasitoid | | Specialist | | Yes | | Flight | | Aerial | | Nocturnal | | No | | 15 | | 20 | 35 | | |
|  | Dryinidae | | Hymenoptera | | Parasitoid | | Specialist | | Yes | | Flight | | Aerial | | Nocturnal | | No | | 0 | | 2 | 2 | | |
|  | Eucharitidae | | Hymenoptera | | Parasitoid | | Specialist | | Yes | | Flight | | Aerial | | Nocturnal | | No | | 5 | | 2 | 7 | | |
|  | Diapriidae | | Hymenoptera | | Parasitoid | | Specialist | | Yes | | Flight | | Aerial | | Nocturnal | | No | | 97 | | 379 | 476 | | |
|  | Encyrtidae | | Hymenoptera | | Parasitoid | | Specialist | | Yes | | Flight | | Aerial | | Nocturnal | | No | | 288 | | 589 | | | 877 |
| Family | | Order | | Forg | | DB | | AS | | DP | | ST | | AP | | OH | | 2017 | | 2018 | | | Sum | |
|  | Eulophidae | | Hymenoptera | | Parasitoid | | Specialist | | Yes | | Flight | | Aerial | | Nocturnal | | No | | 136 | | 222 | | | 358 |
|  | Eupelmidae | | Hymenoptera | | Parasitoid | | Specialist | | Yes | | Flight | | Aerial | | Nocturnal | | No | | 1 | | 1 | | | 2 |
|  | Eurytomidae | | Hymenoptera | | Parasitoid | | Specialist | | Yes | | Flight | | Aerial | | Nocturnal | | No | | 11 | | 11 | | | 22 |
|  | Megaspilidae | | Hymenoptera | | Parasitoid | | Specialist | | Yes | | Flight | | Aerial | | Nocturnal | | No | | 3 | | 10 | | | 13 |
|  | Mymaridae | | Hymenoptera | | Parasitoid | | Specialist | | Yes | | Flight | | Aerial | | Nocturnal | | No | | 22 | | 48 | | | 70 |
|  | Perilampidae | | Hymenoptera | | Parasitoid | | Specialist | | Yes | | Flight | | Aerial | | Nocturnal | | No | | 8 | | 3 | | | 11 |
|  | Platygastridae | | Hymenoptera | | Parasitoid | | Specialist | | Yes | | Flight | | Aerial | | Nocturnal | | No | | 138 | | 161 | | | 299 |
|  | Proctotrupidae | | Hymenoptera | | Parasitoid | | Specialist | | Yes | | Flight | | Aerial | | Nocturnal | | No | | 6 | | 33 | | | 39 |
|  | Pteromalidae | | Hymenoptera | | Parasitoid | | Specialist | | Yes | | Flight | | Aerial | | Nocturnal | | No | | 82 | | 103 | | | 185 |
|  | Signiphoridae | | Hymenoptera | | Parasitoid | | Specialist | | Yes | | Flight | | Aerial | | Nocturnal | | No | | 2 | | 1 | | | 3 |
|  | Torymidae | | Hymenoptera | | Parasitoid | | Specialist | | Yes | | Flight | | Aerial | | Nocturnal | | No | | 2 | | 2 | | | 4 |
|  | Trichogrammatidae | | Hymenoptera | | Parasitoid | | Specialist | | Yes | | Flight | | Aerial | | Nocturnal | | No | | 27 | | 64 | | | 91 |
|  | Braconidae | | Hymenoptera | | Parasitoid | | Specialist | | Yes | | Flight | | Aerial | | Nocturnal | | No | | 158 | | 496 | | | 654 |
|  | Ichneumonidae | | Hymenoptera | | Parasitoid | | Specialist | | Yes | | Flight | | Aerial | | Nocturnal | | No | | 100 | | 238 | | | 338 |
|  | Cynipidae | | Hymenoptera | | Herbivore | | Specialist | | No | | Flight | | Aerial | | Nocturnal | | No | | 135 | | 324 | | | 459 |
|  | Agelenidae | | Araneae | | Carnivore | | Generalist | | No | | Both | | Ground/Veg | | Both | | No | | 1 | | 3 | | | 4 |
|  | Araneidae | | Araneae | | Carnivore | | Generalist | | No | | Both | | Ground/Veg | | Diurnal | | No | | 1 | | 1 | | | 2 |
|  | Clubionidae | | Araneae | | Carnivore | | Generalist | | No | | Ground | | Ground/Veg | | Nocturnal | | No | | 1 | | 0 | | | 1 |
|  | Dictynidae | | Araneae | | Carnivore | | Generalist | | No | | Ground | | Ground/Veg | | Diurnal | | No | | 0 | | 4 | | | 4 |
|  | Mysmenidae | | Araneae | | Carnivore | | Generalist | | No | | Both | | Ground/Veg | | Diurnal | | No | | 1 | | 1 | | | 2 |
|  | Nesticidae | | Araneae | | Carnivore | | Generalist | | No | | Ground | | Ground/Veg | | Diurnal | | No | | 15 | | 19 | | | 34 |
|  | Oxyopidae | | Araneae | | Carnivore | | Generalist | | No | | Both | | Ground/Veg | | Diurnal | | No | | 5 | | 3 | | | 8 |
|  | Selenopidae | | Araneae | | Carnivore | | Generalist | | No | | Ground | | Ground/Veg | | Nocturnal | | No | | 2 | | 0 | | | 2 |
|  | Salticidae | | Araneae | | Carnivore | | Generalist | | No | | Ground | | Ground/Veg | | Diurnal | | No | | 0 | | 2 | | | 2 |
| Family | | Order | | Forg | | DB | | AS | | DP | | ST | | AP | | OH | | 2017 | | 2018 | | | Sum | |
|  | Tetragnathidae | | Araneae | | Carnivore | | Generalist | | No | | Both | | Ground/Veg | | Both | | No | | 6 | | 13 | | | 19 |
|  | Theridiidae | | Araneae | | Carnivore | | Generalist | | No | | Both | | Ground/Veg | | Diurnal | | No | | 1 | | 27 | | | 28 |
|  | Thomisidae | | Araneae | | Carnivore | | Generalist | | No | | Both | | Ground/Veg | | Diurnal | | No | | 0 | | 4 | | | 4 |
|  | Gnaphosidae | | Araneae | | Carnivore | | Generalist | | No | | Ground | | Ground/Veg | | Nocturnal | | No | | 15 | | 13 | | | 28 |
|  | Hahniidae | | Araneae | | Carnivore | | Generalist | | No | | Ground | | Ground/Veg | | Diurnal | | No | | 7 | | 8 | | | 15 |
|  | Lycosidae | | Araneae | | Carnivore | | Generalist | | No | | Ground | | Ground/Veg | | Both | | No | | 89 | | 156 | | | 245 |
|  | Oonopidae | | Araneae | | Carnivore | | Generalist | | No | | Ground | | Ground/Veg | | Nocturnal | | No | | 0 | | 2 | | | 2 |
|  | Phrurolithidae | | Araneae | | Carnivore | | Generalist | | No | | Ground | | Ground/Veg | | Nocturnal | | No | | 0 | | 2 | | | 2 |
|  | Pisauridae | | Araneae | | Carnivore | | Generalist | | No | | Ground | | Ground/Veg | | Nocturnal | | No | | 1 | | 0 | | | 1 |
|  | Titanoecidae | | Araneae | | Carnivore | | Generalist | | No | | Ground | | Ground/Veg | | Nocturnal | | No | | 1 | | 0 | | | 1 |
|  | Plutellidae | | Lepidoptera | | Herbivore | | Specialist | | Yes | | Flight | | Aerial | | Nocturnal | | Crop | | 3202 | | 753 | | | 3955 |
|  | Aphidoidae | | Hemiptera | | Herbivore | | Specialist | | Yes | | Flight | | Ground/Veg | | Diurnal | | Crop | | 32702 | | 8009 | | | 40711 |
|  | Chrysomelidae | | Coleoptera | | Herbivore | | Specialist | | Yes | | Flight | | Aerial | | Diurnal | | Crop | | 361 | | 232 | | | 593 |
|  | Agromyzidae | | Diptera | | Herbivore | | Generalist | | Yes | | Flight | | Aerial | | Diurnal | | Crop | | 838 | | 992 | | | 1830 |
|  | Thripidae | | Thysanoptera | | Herbivore | | Generalist | | Yes | | Flight | | Aerial | | Diurnal | | Crop | | 1379 | | 2032 | | | 3411 |
|  | Tephritidae | | Diptera | | Herbivore | | Generalist | | Yes | | Flight | | Aerial | | Diurnal | | No | | 32 | | 59 | | | 91 |
|  | Brachycera | | Diptera | | Herbivore | | Generalist | | No | | Flight | | Aerial | | Diurnal | | No | | 7020 | | 12765 | | | 19785 |
|  | Nematocera | | Diptera | | Herbivore | | Generalist | | No | | Flight | | Aerial | | Nocturnal | | No | | 13862 | | 34575 | | | 48437 |
|  | Cicadellidae | | Hemiptera | | Herbivore | | Generalist | | No | | Flight | | Aerial | | Diurnal | | No | | 1053 | | 627 | | | 1680 |
|  | Aderidae | | Coleoptera | | Fungivores | | Generalist | | No | | Flight | | Aerial | | Nocturnal | | No | | 0 | | 22 | | | 22 |
|  | Alexiidae | | Coleoptera | | Fungivores | | Specialist | | No | | Flight | | Aerial | | Nocturnal | | No | | 25 | | 13 | | | 38 |
|  | Anobiidae | | Coleoptera | | Herbivore | | Generalist | | No | | Flight | | Aerial | | Diurnal | | No | | 0 | | 3 | | | 3 |
|  | Anthicidae | | Coleoptera | | Omnivore | | Generalist | | No | | Flight | | Aerial | | Diurnal | | No | | 5 | | 11 | | | 16 |
|  | Bruchidae | | Coleoptera | | Herbivore | | Specialist | | Yes | | Flight | | Aerial | | Diurnal | | No | | 2 | | 2 | | | 4 |
| Family | | Order | | Forg | | DB | | AS | | DP | | ST | | AP | | OH | | 2017 | | 2018 | | | Sum | |
|  | Cerylonidae | | Coleoptera | | Fungivores | | Generalist | | No | | Flight | | Aerial | | Nocturnal | | No | | 0 | | 1 | | | 1 |
|  | Corylophidae | | Coleoptera | | Fungivores | | Specialist | | No | | Flight | | Aerial | | Nocturnal | | No | | 3 | | 15 | | | 18 |
|  | Cryptophagidae | | Coleoptera | | Fungivores | | Specialist | | No | | Flight | | Aerial | | Nocturnal | | No | | 107 | | 334 | | | 441 |
|  | Curculionidae | | Coleoptera | | Herbivore | | Generalist | | No | | Flight | | Aerial | | Nocturnal | | No | | 21 | | 15 | | | 36 |
|  | Dermestidae | | Coleoptera | | Herbivore | | Generalist | | No | | Flight | | Aerial | | Diurnal | | No | | 5 | | 0 | | | 5 |
|  | Dytiscidae | | Coleoptera | | Carnivore | | Generalist | | No | | Flight | | Aerial | | Nocturnal | | No | | 2 | | 1 | | | 3 |
|  | Elateridae | | Coleoptera | | Herbivore | | Generalist | | No | | Flight | | Aerial | | Diurnal | | No | | 4 | | 19 | | | 23 |
|  | Eucinetidae | | Coleoptera | | Fungivores | | Specialist | | No | | Flight | | Aerial | | Diurnal | | No | | 2 | | 0 | | | 2 |
|  | Heteroceridae | | Coleoptera | | Carnivore | | Generalist | | No | | Flight | | Aerial | | Diurnal | | No | | 15 | | 0 | | | 15 |
|  | Histeridae | | Coleoptera | | Carnivore | | Generalist | | No | | Flight | | Aerial | | Nocturnal | | No | | 1 | | 0 | | | 1 |
|  | Hydrophilidae | | Coleoptera | | Carnivore | | Generalist | | No | | Flight | | Aerial | | Nocturnal | | No | | 4 | | 0 | | | 4 |
|  | Meloidae | | Coleoptera | | Herbivore | | Generalist | | No | | Flight | | Aerial | | Diurnal | | No | | 1 | | 0 | | | 1 |
|  | Latridiidae | | Coleoptera | | Fungivores | | Specialist | | No | | Flight | | Aerial | | Nocturnal | | No | | 218 | | 413 | | | 631 |
|  | Leiodidae | | Coleoptera | | Fungivores | | Specialist | | No | | Flight | | Aerial | | Nocturnal | | No | | 10 | | 34 | | | 44 |
|  | Micropeplidae | | Coleoptera | | Fungivores | | Specialist | | No | | Flight | | Aerial | | Nocturnal | | No | | 4 | | 1 | | | 5 |
|  | Monotomidae | | Coleoptera | | Fungivores | | Specialist | | No | | Flight | | Aerial | | Nocturnal | | No | | 0 | | 1 | | | 1 |
|  | Mordellidae | | Coleoptera | | Herbivore | | Generalist | | No | | Flight | | Aerial | | Diurnal | | No | | 2 | | 3 | | | 5 |
|  | Mycetophagidae | | Coleoptera | | Fungivores | | Specialist | | No | | Flight | | Aerial | | Nocturnal | | No | | 58 | | 105 | | | 163 |
|  | Nitidulidae | | Coleoptera | | Fungivores | | Specialist | | No | | Flight | | Aerial | | Diurnal | | No | | 66 | | 165 | | | 231 |
|  | Prostomidae | | Coleoptera | | Fungivores | | Generalist | | No | | Flight | | Aerial | | Nocturnal | | No | | 1 | | 0 | | | 1 |
|  | Pselaphidae | | Coleoptera | | Carnivore | | Generalist | | No | | Flight | | Aerial | | Nocturnal | | No | | 1 | | 4 | | | 5 |
|  | Ptiliidae | | Coleoptera | | Fungivores | | Specialist | | No | | Flight | | Aerial | | Nocturnal | | No | | 10 | | 0 | | | 10 |
|  | Scarabaeidae | | Coleoptera | | Herbivore | | Generalist | | No | | Flight | | Aerial | | Nocturnal | | No | | 1 | | 0 | | | 1 |
|  | Scolytidae | | Coleoptera | | Herbivore | | Generalist | | No | | Flight | | Aerial | | Nocturnal | | No | | 168 | | 141 | | | 309 |
| Family | | Order | | Forg | | DB | | AS | | DP | | ST | | AP | | OH | | 2017 | | 2018 | | | Sum | |
|  | Scydmaenidae | | Coleoptera | | Carnivore | | Specialist | | No | | Flight | | Aerial | | Nocturnal | | No | | 0 | | 1 | | | 1 |
|  | Silvanidae | | Coleoptera | | Herbivore | | Specialist | | No | | Flight | | Aerial | | Nocturnal | | No | | 12 | | 48 | | | 60 |
|  | Silphidae | | Coleoptera | | Omnivore | | Generalist | | No | | Flight | | Aerial | | Nocturnal | | No | | 0 | | 1 | | | 1 |
|  | Tenebrionidae | | Coleoptera | | Herbivore | | Generalist | | No | | Flight | | Aerial | | Diurnal | | No | | 0 | | 1 | | | 1 |
|  | Apidae | | Hymenoptera | | Herbivore | | Generalist | | No | | Flight | | Aerial | | Diurnal | | No | | 258 | | 200 | | | 458 |
|  | Tenthredinidae | | Hymenoptera | | Herbivore | | Generalist | | No | | Flight | | Aerial | | Diurnal | | No | | 18 | | 35 | | | 53 |
|  | Scoliidae | | Hymenoptera | | Carnivore | | Generalist | | No | | Flight | | Aerial | | Diurnal | | No | | 1 | | 1 | | | 2 |
|  | Pompilidae | | Hymenoptera | | Carnivore | | Specialist | | No | | Flight | | Aerial | | Diurnal | | No | | 0 | | 10 | | | 10 |
|  | Psocidae | | Psocoptera | | Fungivores | | Generalist | | No | | Flight | | Aerial | | Diurnal | | No | | 52 | | 33 | | | 85 |
|  | Cydnidae | | Hemiptera | | Herbivore | | Generalist | | No | | Flight | | Ground/Veg | | Diurnal | | No | | 1 | | 15 | | | 16 |
|  | Miridae | | Hemiptera | | Herbivore | | Generalist | | No | | Flight | | Aerial | | Diurnal | | No | | 118 | | 272 | | | 390 |
|  | Tingidae | | Hemiptera | | Herbivore | | Generalist | | No | | Flight | | Aerial | | Diurnal | | No | | 2 | | 14 | | | 16 |
|  | Plataspidae | | Hemiptera | | Herbivore | | Generalist | | No | | Flight | | Aerial | | Diurnal | | No | | 4 | | 1 | | | 5 |
|  | Berytidae | | Hemiptera | | Herbivore | | Generalist | | No | | Flight | | Aerial | | Diurnal | | No | | 1 | | 0 | | | 1 |
|  | Lygaeidae | | Hemiptera | | Herbivore | | Generalist | | No | | Flight | | Aerial | | Diurnal | | No | | 650 | | 303 | | | 953 |
|  | Pentatomidae | | Hemiptera | | Herbivore | | Generalist | | No | | Flight | | Aerial | | Diurnal | | No | | 13 | | 2 | | | 15 |
|  | Coreidae | | Hemiptera | | Herbivore | | Generalist | | No | | Flight | | Aerial | | Diurnal | | No | | 8 | | 8 | | | 16 |
|  | Tessaratomidae | | Hemiptera | | Herbivore | | Generalist | | No | | Flight | | Aerial | | Diurnal | | No | | 0 | | 2 | | | 2 |
|  | Rhyparochromidae | | Hemiptera | | Herbivore | | Generalist | | No | | Flight | | Aerial | | Diurnal | | No | | 3 | | 0 | | | 3 |
|  | Urostylidae | | Hemiptera | | Herbivore | | Generalist | | No | | Flight | | Aerial | | Diurnal | | No | | 0 | | 7 | | | 7 |
|  | Pyrrhocoridae | | Hemiptera | | Herbivore | | Generalist | | No | | Flight | | Aerial | | Diurnal | | No | | 2 | | 0 | | | 2 |
|  | Scutelleridae | | Hemiptera | | Herbivore | | Generalist | | No | | Flight | | Aerial | | Diurnal | | No | | 2 | | 0 | | | 2 |
|  | Termitidae | | Blattodea | | Herbivore | | Generalist | | No | | Flight | | Aerial | | Nocturnal | | No | | 0 | | 3 | | | 3 |
|  | Crambidae | | Lepidoptera | | Herbivore | | Generalist | | No | | Flight | | Aerial | | Nocturnal | | No | | 33 | | 1 | | | 34 |
| Family | | Order | | Forg | | DB | | AS | | DP | | ST | | AP | | OH | | 2017 | | 2018 | | | Sum | |
|  | Membracidae | | Hemiptera | | Herbivore | | Specialist | | No | | Flight | | Aerial | | Diurnal | | No | | 17 | | 17 | | | 34 |
|  | Ephemeridae | | Ephemeroptera | | Herbivore | | Generalist | | No | | Flight | | Aerial | | Diurnal | | No | | 6 | | 13 | | | 19 |
|  | Hesperiidae | | Lepidoptera | | Herbivore | | Generalist | | No | | Flight | | Aerial | | Diurnal | | No | | 9 | | 9 | | | 18 |
|  | Psyllidae | | Hemiptera | | Herbivore | | Specialist | | No | | Flight | | Aerial | | Diurnal | | No | | 28 | | 252 | | | 280 |
|  | Cercopidae | | Hemiptera | | Herbivore | | Generalist | | No | | Flight | | Aerial | | Diurnal | | No | | 0 | | 23 | | | 23 |
|  | Nymphalidae | | Lepidoptera | | Herbivore | | Generalist | | No | | Flight | | Aerial | | Diurnal | | No | | 1 | | 2 | | | 3 |
|  | Tridactylidae | | Orthoptera | | Omnivore | | Generalist | | No | | Ground | | Ground/Veg | | Diurnal | | No | | 0 | | 3 | | | 3 |
|  | Gryllotalpidae | | Orthoptera | | Herbivore | | Generalist | | No | | Flight | | Aerial | | Nocturnal | | No | | 2 | | 4 | | | 6 |
|  | Gryllidae | | Orthoptera | | Herbivore | | Generalist | | No | | Flight | | Aerial | | Nocturnal | | No | | 26 | | 110 | | | 136 |
|  | Coenagrionidae | | Odonata | | Carnivore | | Generalist | | No | | Flight | | Aerial | | Diurnal | | No | | 1 | | 0 | | | 1 |
|  | Ectobiidae | | Blattodea | | Omnivore | | Generalist | | No | | Flight | | Aerial | | Nocturnal | | No | | 0 | | 93 | | | 93 |
|  | Centipede | | Scolopendromopha | | Carnivore | | Generalist | | No | | Ground | | Ground/Veg | | Nocturnal | | No | | 47 | | 57 | | | 104 |
