## Supplementary material for "Contrasting roles of landscape compositions in shaping functional traits of arthropod community in subtropical vegetable fields": able S2

**Table S2. Coefficients and SE from model averaging showing the relationship between functional diversity index of arthropod community and landscape variables and interactions with sampling regions at a 100, 200, 300, 400 and 500 m radius**. A plus (**+**) indicates that a significant or interaction with sampling regions, and asterisks (*) show significance levels (*≤0.05; **≤0.01; ***<0.001).

| **Response** | **Scale**  **(m)** | **AICc** | **Intercept** | **Region** | **Non-cruciferous** | **Forest** | **Grassland** | **Urban** | **Water** | **Interaction with sampling regions** | | | | |
| --- | --- | --- | --- | --- | --- | --- | --- | --- | --- | --- | --- | --- | --- | --- |
|  |  |  |  |  |  |  |  |  |  | **Non-cruciferous** | **Forest** | **Grassland** | **Urban** | **Water** |
| FRic | 100 | 231.2 | 3.06±0.06*** | **-** | -0.01±0.02 | **0.12±0.04**** | 0.06±0.06 | - | 0.06±0.04 | **-** | **-** | **-** | **-** | **-** |
|  | 200 | 227.0 | 3.04±0.06*** | **-** | **-** | **0.12±0.03**** | **-** | 0.01±0.03 | **-** | **-** | **-** | **-** | **-** | **-** |
|  | 300 | 229.0 | 3.04±0.06*** | **-** | **-** | **0.11±0.03***** | **-** | 0.01±0.02 | **-** | **-** | **-** | **-** | **-** | **-** |
|  | 400 | 230.3 | 3.04±0.05*** | **-** | **-** | **0.11±0.03**** | **-** | 0.02±0.03 | **-** | **-** | **-** | **-** | **-** | **-** |
|  | 500 | 227.0 | 3.04±0.06*** | **+** | **-** | **0.13±0.03**** | **-** | 0.05±0.04 | -0.07±0.12 | **-** | **-** | **-** | **-** | **+** |
| FEve | 100 | -188.7 | -1.29±0.04*** | **+** | **-0.08±0.04*** | **-** | **-** | **-** | **-** | **-** | **-** | **-** | **-** | **-** |
|  | 200 | -191.0 | -1.29±0.03*** | **+** | - | 0.01±0.02 | 0.04±0.04 | 0.06±0.04 | **-** | **-** | **-** | **-** | **-** | **-** |
|  | 300 | -193.2 | -1.33±0.03*** | **-** | -0.01±0.01 | 0.01±0.01 | **0.09±0.03**** | 0.03±0.04 | -0.02±0.03 | **-** | **-** | **-** | **-** | **-** |
|  | 400 | -196.6 | -1.33±0.03*** | **-** | -0.01±0.03 | **-** | **0.10±0.03***** | **-** | -0.08±0.03 | **-** | **-** | **-** | **-** | **-** |
|  | 500 | -192.1 | -1.33±0.03*** | **-** | **-** | **-** | **0.08±0.03**** | **-** | **-0.09±0.04*** | **-** | **-** | **-** | **-** | **-** |
| FDis | 100 | -139.4 | -1.36±0.11*** | **-** | 0.01±0.01 | **-** | -0.02±0.04 | **-** | **-** | - | - | - | - | **-** |
|  | 200 | -141.3 | -1.35±0.11*** | **-** | **-** | **-** | 0.08±0.06 | **-** | **-** | - | - | - | - | **-** |
|  | 300 | -144.4 | -1.36±0.11*** | **-** | **-** | **-** | **0.13±0.05**** | **-** | **-** | - | - | - | - | **-** |
|  | 400 | -144.3 | -1.36±0.12*** | **-** | **-** | 0.01±0.03 | **0.13±0.05**** | **-** | **-** | - | - | - | - | **-** |
|  | 500 | -142.3 | -1.36±0.12*** | **-** | **-** | 0.02±0.04 | **0.12±0.05*** | **-** | **-** | - | - | - | - | **-** |
