## Supplementary figures and images for "Contrasting roles of landscape compositions in shaping functional traits of arthropod community in subtropical vegetable fields"

### Figure S1

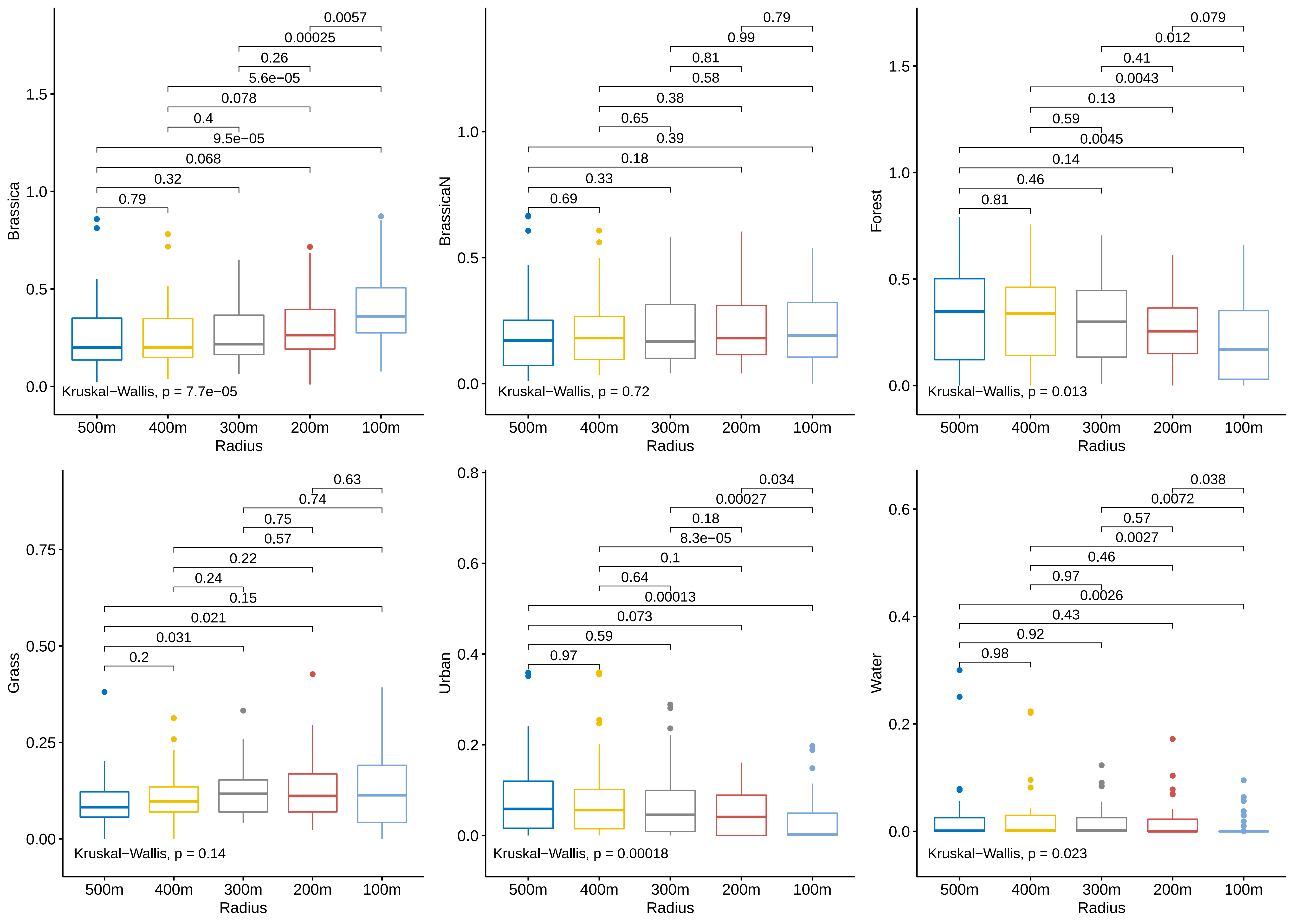
